# Type 4 pili mediated natural competence in *Fusobacterium nucleatum*

**DOI:** 10.1101/2023.06.09.544391

**Authors:** Blake E. Sanders, Ariana Umana, Tam T.D. Nguyen, Kevin J. Williams, Christopher C. Yoo, Michael A. Casasanta, Bryce Wozniak, Daniel J. Slade

## Abstract

Many bacterial species naturally take up DNA from their surroundings and recombine it into their chromosome through homologous gene transfer (HGT) to aid in survival and gain advantageous functions. Herein we present the first characterization of Type 4 pili mediated natural competence in *Fusobacterium nucleatum*, which are Gram-negative, anaerobic bacteria that participate in a range of infections and diseases including periodontitis, preterm birth, and cancer. We bioinformatically identified components of the Type 4 conjugal pilus machinery and show this is a conserved system within the *Fusobacterium* genus. We next validate Type 4 pili in natural competence in *F. nucleatum* strain 23726 and show that gene deletions in key components of pilus deployment (*pilQ)* and cytoplasmic DNA import (*comEC)* abolish DNA uptake and chromosomal incorporation. We next show that natural competence may require native *F. nucleatum* DNA methylation to bypass restriction modification systems and allow subsequent genomic homologous recombination. In summary, this proof of principle study provides the first characterization of natural competence in *Fusobacterium nucleatum* and highlights the potential to exploit this DNA import mechanism as a genetic tool to characterize virulence mechanisms of an opportunistic oral pathogen.

## INTRODUCTION

A striking feature of bacteria is their ability to be genomic ‘shape-shifters’ capable of gaining and losing genes in a systematic and regulated manner, which can drive evolution, adaptation to surroundings, and virulence^1,2^. Unfortunately, these methods also provide bacteria a way to acquire antibiotic resistance genes; an ever-increasing problem associated with human pathogens^3^. Bacteria naturally accept DNA using three mechanisms: conjugation, transduction (viral), and natural competence (NC)^4^. NC is a method in which bacteria import DNA from their surroundings through the deployment of a surface exposed protein complex known as the Type 4 competence pilus. Although NC is an important component of bacterial adaptation in native settings, it is often difficult to initiate in laboratory settings and to date only ∼80 different strains of bacteria have been confirmed to be naturally competent^5,6^. However, once harnessed, bacterial NC has proven to be an efficient means of genetic manipulation in the laboratory^7^; especially in situations where the target bacterium has proven recalcitrant to other methods of DNA introduction including chemotransformation, electroporation, sonoporation^8^, and conjugational transfer^9^.

NC encompasses three major stages: (i) DNA uptake, (ii) recombination of homologous DNA onto the chromosome, and (iii) expression of acquired genetic material. DNA uptake comprises the binding of external DNA to the pilus through pilin proteins, followed by import of the DNA through a ratcheting mechanism that guides the DNA through the outer membrane and into the periplasm in Gram-negative bacteria where it is unwound into single-stranded DNA (ssDNA) while being delivered to the cytoplasmic membrane^10^. The second step of NC involves the translocation of ssDNA across the cytoplasmic membrane. This process is highly conserved among both Gram-negative and Gram-positive bacteria and is mediated by a cytoplasmic membrane channel (ComEC/ComA) and other cytoplasmic proteins found in all known competent bacteria^11^.

A barrier to efficient transformation of exogenous DNA is the presence of restriction modification (R-M) systems which consist of strain specific DNA methyltransferases (DMTase) that mark DNA as ‘self’, as well as nucleases that recognize and cleave entering DNA that has non-native methylation patterns, which is considered foreign and a threat to genome integrity^12^. These systems have previously been overcome by using native DMTases to imitate the methylation pattern of the host bacteria, thereby bypassing the R-M system nucleases and continuing on to incorporation into the genome^13^. This approach was recently utilized by our lab to greatly enhance gene knockout and complementation in *F. nucleatum*^14^. Previous studies have shown that not only must the imported DNA have high homology to the accepting organism for chromosomal incorporation through homologous recombination, but it may also require specific DNA methylation patterns for recognition and import^15^, as well as protection from nucleases during and after chromosomal incorporation^16-18^.

Prior to the work presented here, there has been only one report mentioning any prediction of Type 4 pili in *Fusobacterium* by Desvaux et al, which reported the initial identification and investigation into Type 4 pili in *Fusobacterium* through a bioinformatic analysis of the protein secretion systems in *F. nucleatum* subsp. *nucleatum* 25586 (*F. nucleatum* 25586)^19^. Our bioinformatic analysis expanded upon this investigation to identify a full repertoire of Type 4 pili assembly genes present in multiple *Fusobacterium species*. We next set out to determine if *Fusobacterium* are naturally competent, with a focus on *F. nucleatum* subsp. *nucleatum* 23726 (*F. nucleatum* 23726). Our results show that *F. nucleatum* 23726 is naturally competent and that using native DMTase modification of genomic and plasmid DNA is critical for transformation and incorporation of genetic elements onto the chromosome. We show that this process is Type 4 pili dependent as knocking out genes critical for deployment of the pilus through the outer membrane (*pilQ*) and import of DNA into the cytoplasm *(comEC*) abolishes NC. We show that both chromosomal and plasmid DNA is imported by pili and upon further development and investigation could lead to a new method for genetic manipulation in *F. nucleatum*.

## RESULTS

### Bioinformatic analysis reveals conservation of natural competence genes in *Fusobacterium*

Previous genomic analyses of *F. nucleatum* 25586 revealed some of the genes encoding the machinery required for the Type 4 pilin/fimbriation secretion system including homologs of PilD, PilC, PilB, PilQ, and PilT^19^. However, questions remained on whether these genes belonged to a Type 4 protein secretion system, Type 4 pili mediated natural competence system, or a Type 2 protein secretion system. Using *Fusobacterium* genomes present in the FusoPortal database^20,21^, we uncovered a repertoire of genes associated with the Type 4 pilin/fimbriation system (**Fig. 1A,B**), and we note their genome location in *F. nucleatum* 23726 (**Fig. 1C**). Interestingly, this appears to be a minimalist system that is missing genes that were deemed necessary for the natural competence machinery in other bacteria. For example, the five known components of the Type IV pilus competency machinery that were lacking in *F. nucleatum* include PilF, an outer membrane lipoprotein essential for biogenesis and localization of the secretin, and PilM, PilN, PilO, and PilP; four proteins found to make up the inner membrane pilus subcomplex. The functional roles of the inner membrane associated proteins of the Type IV pilus are not as well defined as other components of the system. According to co-fractionation data and protein stability phenotypes, PilM, PilN, PilO, and PilP are predicted to form an inner membrane subcomplex that could be involved in aligning the outer membrane secretin with the pilus assembly machinery^22^. In the well-studied Gram-negative pathogen, *Pseudomonas aeruginosa*, PilP is suggested to bridge the inner and outer membrane complexes, forming a continuous channel across the periplasm through which the pilus can efficiently extend and retract^23^. This arrangement would guarantee that the pore formed by PilQ, the secretin, is aligned with the inner membrane complex and the cytoplasmic ATPases involved in pilus movement^23^. Although *F. nucleatum* lacks these important structural components, we show below that *F. nucleatum* Type IV pili-mediated NC does not require these proteins for transfer of exogenous DNA. Examination of the other proteins of this seemingly novel and minimalistic competence machinery is warranted to understand their functions in *Fusobacterium* horizontal gene transfer.

**Figure 1.**
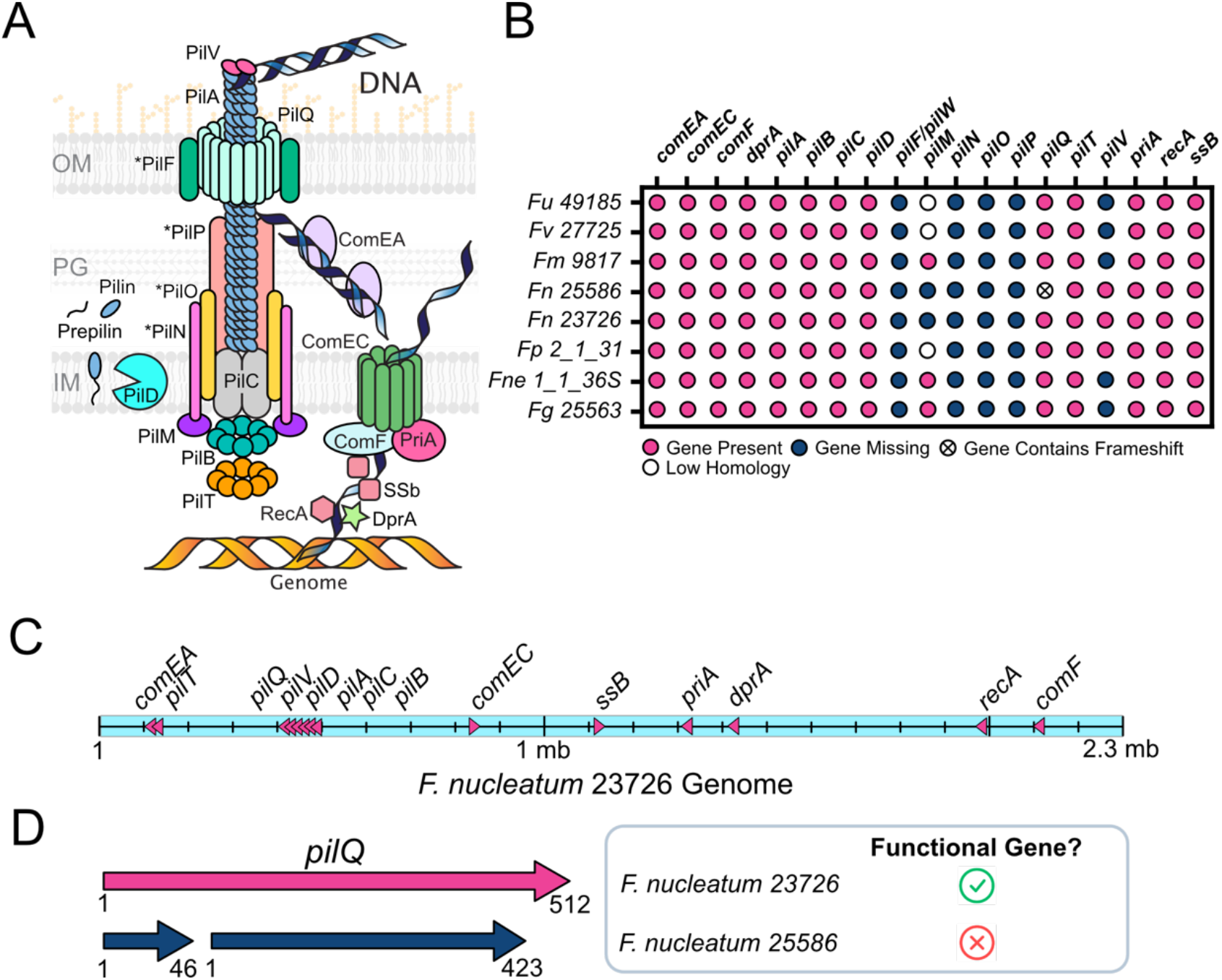
Bioinformatic analysis of *Fusobacterium* Type 4 pili components. **(A)** Schematic of Type 4 pili proteins identified in *F. nucleatum* and their proposed assembly based on the characterization of Type 4 pili in other bacteria. * Stars on proteins in the schematic indicate that *Fusobacterium* do not contain these genes. **(B)** Identification of Type 4 pilus genes in seven species of *Fusobacterium* that covers 8 strains. **(C)** Chromosomal location of pilus genes in the strain *F. nucleatum* subsp. *nucleatum* 23736. **(D)** Analysis of the *pilQ* genes in *F. nucleatum* 23726 and *F. nucleatum* 25586, showing a frame shift creating an inactive pilus protein in *F. nucleatum* 25586. *Fu: Fusobacterium ulcerans; Fv: Fusobacterium varium; Fm: Fusobacterium mortiferum; Fn: Fusobacterium nucleatum; Fp: Fusobacterium periodonticum; Fne: Fusobacterium necrophorum; Fg: Fusobacterium gonidiaformans*.

Next, we discovered that the *pilQ* gene in *F. nucleatum* 25586 contains a frameshift that divides the gene, which would make it unable to deploy pili for DNA import (**Fig. 1D**), rendering this strain naturally incompetent as we highlight experimentally. Genes highlighted in **Figure 1B** are detailed in **Table S1**.

### Type 4 pili mediated import and chromosomal incorporation of *F. nucleatum* genomic DNA

Our initial experiment was to determine if chromosomal DNA marked with the chloramphenicol resistance gene *catP* (*F. nucleatum* 23726 *ΔgalKT fadA::FLAG-fadA* (Cm^r^) could be imported by wild-type *F. nucleatum* 23726 and *F. nucleatum* 23726 *ΔgalKT* and be incorporated into the genome as determined by the presence of new thiamphenicol resistant *F. nucleatum* 23726 (**Fig. 2A**)^24^. As shown in **Figure 2B**, incubating 5 μg of antibiotically marked chromosomal DNA results in colonies for both *F. nucleatum* 23726 and *F. nucleatum* 23726 *ΔgalKT*, but not strains that lack *pilQ* (*F. nucleatum* 23726 *ΔgalKT pilQ*) and *comEC* (*F. nucleatum* 23726 *ΔgalKT comEC*). Details of the gene deletion method for *pilQ* and *comEC* are provided in detail in **Figure S1**.

**Figure 2.**
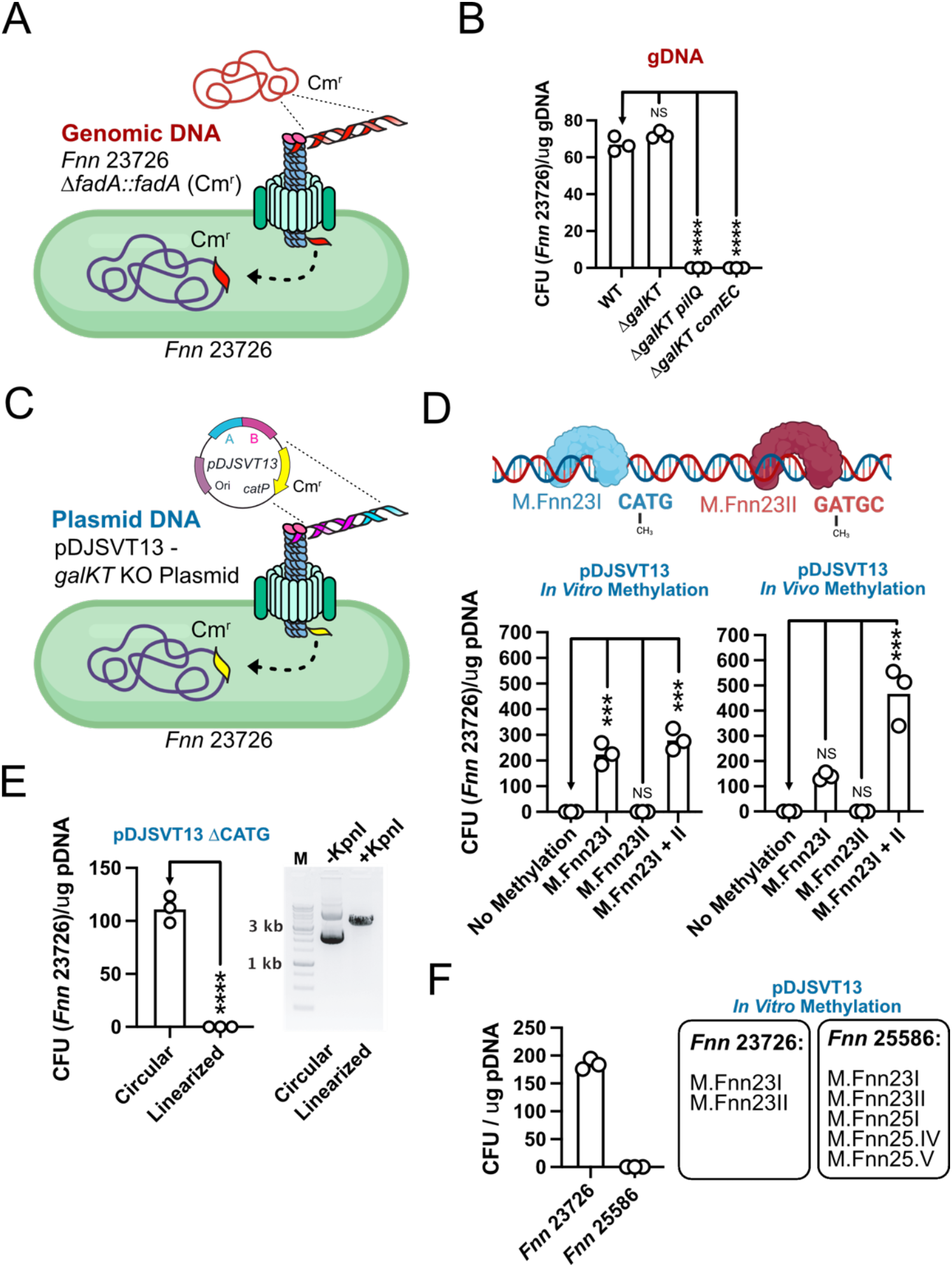
*Fusobacterium nucleatum* utilize Type 4 pili for natural competence of genomic and plasmid DNA. **(A)** Schematic of genomic DNA uptake of a chloramphenicol marked chromosome from *F. nucleatum* 23726 taken up by the same wild-type strain. **(B)** *F. nucleatum* 23726 can import and chromosomally incorporate exogenous genomic DNA from the same species. Mutations in *pilQ* and *comEC* abolish natural competence. **(C)** Schematic of plasmid DNA uptake and chromosomal incorporation by *F. nucleatum* 23726, resulting in chloramphenicol resistant bacteria. **(D)** Plasmid DNA incorporation onto the chromosome of *F. nucleatum* 23726 is DNA methylation dependent and works by methylating plasmids *in vitro* (purified enzymes M.Fnn23I, M.Fnn23II) and *in vivo* (*E. coli* expressing M.Fnn23I, M.Fnn23II, or both). **(E)** *F. nucleatum* 23726 are naturally competent of the plasmid pDJSVT13 lacking the sequence CATG, which is methylated by M.Fnn23I. Linearization of plasmid abolishes import and/or chromosomal incorporation. **(F)** *F. nucleatum* 25586 is not naturally competent for plasmid DNA (truncated *pilQ* gene), even after treating with five native DNA methyltransferases that were previously used to enable transformation by electroporation. Statistical values are as follows: ns (not significant), P.0.05;*, P,0.05;**, P,0.01;***, P,0.001;****, P,0.0001. For panels B and D data, we used a two-way ANOVA. Panel E used a one-way ANOVA.

These experiments were first tested with exponential phase bacteria (OD_600_=0.5), which serendipitously proved to be the optimal OD_600_ for natural competence in *F. nucleatum* 23726. We next optimized natural competence parameters for *F. nucleatum* 23726 (**Fig. S2**) and show that bacteria grown to an OD_600_=0.5 that are incubated with 5 μg of DNA for 4-hours produces the highest efficiency for chromosomal incorporation. These parameters were used for all experiments in this manuscript unless otherwise specified. Worth noting, we did try overnight, stationary phase bacteria for natural competence studies, but did not find this method successful. We highlight in the discussion that we did not exhaust all options for growth conditions and other factors to test natural competence that have proven successful in other bacteria.

### Native *F. nucleatum* methylation is critical for transformation of plasmid DNA

Initial NC assays incubated with cloning shuttle vector plasmid DNA (pDJSVT13, homologous regions surrounding the genes *galK* and *galT*) were unsuccessful in uptake and integration (**Fig. 2C,D**). Since our initial experiment of using genomic DNA for natural competence was successful, and knowing genomic DNA contains native methylation patterns, we hypothesized that DNA import or protection from nuclease digestion could be methylation dependent. Therefore, we methylated pDJSVT13 with the *F. nucleatum* DMTases M.Fnn23I (CA^m^TG) and M.Fnn23II (GA^m^TGC) and show that this enables natural competence and incorporation of this plasmid onto the chromosome (**Fig 2D**)^14^. As some bacteria require a specific sequence to be methylated for DNA recognition, binding, and uptake (*Campylobacter jejuni:* RAATTY sequence)^15^, we deleted the key methylation sequence (CA^m^TG) from pDJSVT13 to determine if this sequence was needed for uptake. We prove that this sequence is not required for uptake as the pDJSVT13 ΔCATG plasmid, without *F. nucleatum* methylation is imported and chromosomally integrated (**Fig. 2E**). As many bacteria can take up linear DNA (Linearized plasmid, PCR products), we linearized pDJSVT13 ΔCATG and show that this abolishes either DNA uptake and/or chromosomal integration. Thus, we hypothesize that the role of DNA methyltransferases in *Fusobacterium*, as with many other bacteria, is to protect DNA to circumvent R-M systems in homologous strains and not for pilus recognition and DNA import.

To round out the study, we show that *F. nucleatum* 25586, which contains a frameshift in *pilQ* (**Fig. 1D**), is not naturally competent (**Fig 2F**). This experiment methylated pDJSVT13 with five *F. nucleatum* DNA methyltransferases (M.Fnn23I, M.Fnn23II, M.Fnn25I, M.Fnn25IV, M.Fnn25V) that were previously shown to be necessary for plasmid protection during electroporation and development of a genetic system in *F. nucleatum 25586*^14,24^. Nevertheless, this did not make *F. nucleatum* 25586 take up the DNA, supporting our hypothesis that this strain is not naturally competent because it cannot deploy a functional pilus to bind and import DNA in the absence of the outer membrane channel formed by PilQ.

## DISCUSSION

In this study we show that *Fusobacterium nucleatum* uses Type IV pili mediated natural competence for DNA import and horizontal gene transfer. We bioinformatically identified components of the DNA uptake machinery and discovered that *F. nucleatum* 23726 was able to take up genomic DNA containing a *catP* gene (chloramphenicol resistance), and subsequently induce horizontally transfer of this gene to the chromosome of the wild-type strain. We next show that native DNA methylation is critical, potentially even necessary, to bypass R-M system cleavage of entering DNA during homologous recombination onto the chromosome. We finally show that DNA methylation is not necessary for pilus recognition and DNA import as deletion of the dominant methylation sequence (CA^m^TG) from our knockout plasmid allowed horizontal gene transfer in the absence of DNA methylation. As the importance of DNA methylation for NC was previously reported for *Helicobacter pylori, Pseudomonas stutzeri, C. jejuni, Neisseria meningitis*, and most recently the ESKAPE pathogen *Acinetobacter baumannii*^*25*^, this could indicate that horizontal gene transfer in *Fusobacterium* will likely only occur between bacteria that share similar methylation patterns, making epigenomic homology a key future direction to study to utilize natural competence for molecular genetic applications across *Fusobacterium* species.

To prove this natural competence was Type 4 pili mediated, we show that the pilus proteins PilQ and ComEC are essential for import and translocation of DNA as gene deletions rendered these strains incapable of DNA import. Since the Type 4 pilus in *Fusobacterium* appears to be a minimalist system that lacks key proteins involved in pilus formation, it presents a future direction to study the mechanisms of natural competence in an emerging ‘oncomicrobe’. Interestingly, as *Fusobacterium* is a non-motile bacterium^26^, the lack of the PilMNOPQ complex within the Type 4 pilus could explain why *Fusobacterium* do not display twitching motility, which is attributed to these proteins in *Pseudomonas aeruginosa*^22^. Another component of the *Fusobacterium* Type IV pilus system that is of interest is the minor pilin subunit, PilV. Our bioinformatic analysis revealed that only *F. nucleatum* 23726, *F. nucleatum* 25586, and *F. periodonticum* 2_1_31 contained the minor pilin subunit. Minor pilins have been shown to play crucial roles in the uptake process of exogenous DNA. The minor pilin of *N. meningitidis*, ComP, has been shown to mediate DNA binding to pili, by recognizing a specific DNA uptake sequence (DUS)^27^. In addition, recent structural studies of *T. thermophilus* minor pilin, ComZ, revealed two major domains, where one showed involvement in DNA binding^28^. Future studies characterizing strains of *Fusobacterium* that lack minor pilin genes will be critical to determine the role of minor pilins in DNA uptake during NC.

The repertoire of NC genes are often not constitutively expressed but rather only switched on during certain growth phases or in response to environmental stimuli^25^. Known inducers of competence in other bacteria include high cell density, antibiotic stress, DNA damage, and starvation^25^. We show that *F. nucleatum* NC to be most efficient during exponential phase growth. However, future studies investigating induction of NC in *F. nucleatum* during different growth phases, including adding DNA directly to colonies on a plate, could be beneficial for increasing NC efficiency if the development of molecular genetic tools is desired. Finally, considering the vast differences in *Fusobacterium* genotypes and phenotypes, it may be that each species, subspecies, or even strains of bacteria could have specific requirements for the induction of natural competence.

In conclusion, we report the first characterization of Type 4 pili mediated natural competence in *F. nucleatum* and show conservation of this system among the *Fusobacterium* genus. This proof of principle study provides a starting foundation for further exploitation of NC as genetic tool for investigating the role of virulence in *F. nucleatum*.

## MATERIALS AND METHODS

### Bacterial Strains and Growth Conditions

*F. nucleatum* strains ATCC 23726 and 25586 were cultured on solid agar plates made with Columbia Broth (Gibco) substituted with hemin (5 μg/mL) and menadione (0.5 μg/mL) (CBHK) under anaerobic conditions (90% N_2_, 5% H_2_, 5% CO_2_) at 37 °C. Liquid cultures started from single colonies were grown in CBHK media under the same conditions. For plasmid DNA production, *E. coli* strains were grown aerobically overnight at 37 °C in LB (15 g/L NaCl, 15 g/L tryptone, 10 g/L yeast extract). Antibiotics were used where appropriate in the following concentrations: chloramphenicol 10μg/mL, carbenicillin 100 μg/mL, thiamphenicol 5 μg/mL (CBHK plates) and 2.5 μg/mL (CBHK broth). All bacteria strains and plasmids used in these studies are presented in **Table S2** and **S4**, respectively.

### Construction of Δ*pilQ* and Δ*comEC* Mutants

A galactose-selectable gene deletion system developed in our lab was utilized to create markerless gene knockouts of *pilQ* and *comEC* (**Fig. S1**). Briefly, 750 bp directly upstream and downstream of *pilQ* and *comEC* were amplified by PCR, making complementary fragments fused by OLE-PCR. This product was ligated into a cloning shuttle vector using KpnI/MluI restriction sites. This vector was then electroporated (2.5 kV, 50 μF capacitance, 360 OHMS resistance) into competent *F. nucleatum* 23726 Δ*galKT* and selected on thiamphenicol (single-crossover homologous recombination), followed by selection on solid media containing 3% galactose which produces either complete gene deletions or wild-type bacteria revertants through double-crossover homologous recombination. Gene deletions were verified by PCR, RT-PCR, and sequencing (**Figure S1**). All primers were ordered from IDT DNA (**Table S3**).

### RNA Extraction and RT-PCR

*F. nucleatum* cultures were grown to stationary phase and pelleted by high-speed centrifugation (12,000 x g, for 3 minutes at room temperature). TRIzol Extraction Isolation of total RNA was performed following manufacturer’s instructions (Invitrogen). Briefly, cell pellets were resuspended in 1 mL of TRIzol reagent (Invitrogen) and 0.2 mL of chloroform was added. Solution was centrifuged for 15 minutes at 12,000 x g, at 4 °C. The RNA-containing aqueous phase was collected, and the RNA precipitated after 500 μL of isopropanol had been added. The RNA pellet was then washed with 75% ethanol and centrifuged at 10,000 x g, for 5 minutes at 4°C. After drying at room temperature for 10 minutes, the RNA pellet was resuspended in 30 μL sterile RNAse-free water and solubilized by incubating in a water bath at 55 °C for 10 minutes. Total RNA was quantified using Qubit™ RNA HS Assay Kit (ThermoFisher). Before reverse transcriptase (RT)-PCR, RNA samples were subjected to DNAse treatment. Briefly, 500 ng of total RNA was incubated with DNAse I (Invitrogen) for 2 hours at 37 °C. Following treatment, DNAse I was inactivated using EDTA and heating mixture for 5 minutes at 65 °C. (RT)-PCR was performed using the Takara PrimeScript™ One Step RT-PCR Kit according to the manufacturer’s instructions. The PCR conditions consisted of reverse transcription for 30 minutes at 50°C, initial denaturation for 2 minutes at 94 °C, followed by 30 cycles (30 sec at 94 °C, 30 sec at 50-62 °C, and 30 sec at 68 °C) and elongation at 68 °C for 1 minutes. The expected bands around 250 bp was confirmed on a 1.5% agarose gel. Specific primers to detect knockout of gene and validate intact genes upstream and downstream of the gene of interest (**Table S3**) were used to amplify from RNA extracts.

### DNA extraction

Genomic DNA was purified from 3 mL of stationary phase *F. nucleatum* Δ*fadA::fadA* Cm^r^ cultures using Wizard® Genomic DNA Purification Kit (Promega) and quantified using a Nanodrop spectrophotometer. Plasmid pDJSVT13 (*galKT* gene deletion plasmid) was purified from overnight co-expression of in *E. coli* TOP10 cells containing pDJSVT26 that expresses the Type II methyltransferases M.Fnn23I and M.Fnn23II. Both methylated plasmids were purified together using EZ-10 Spin Column Plasmid Miniprep Kit (BioBasic). The pDJSVT26 plasmid, which confers ampicillin resistance, was not separated from pDJSVT13 before adding to *F. nucleatum* 23726 for NC assays.

### *In vitro* plasmid methylation with M.Fnn25I, M.Fnn25IV, and M.Fnn25V

For studies involving the strain *F. nucleatum* 25586, pDJSVT13, which was previously methylated *in vivo* in *E. coli* by enzymes M.Fnn23I and M.Fnn23II, was incubated with 1 μM each of purified M.Fnn25I, M.Fnn25IV, and M.Fnn25V for two hours at 37 °C in CutSmart buffer (NEB) containing 160 μM S-adenosylmethionine (SAM: NEB). This five-enzyme methylation cocktail was previously necessary to achieve genetic alteration in the strain as described in Umaña et al^14^.

### Linearization of plasmid DNA

Miniprep plasmid DNA was digested with restriction enzyme KpnI following manufacturer’s instructions (New England Biolabs). After two hours at 37 °C, miniprep plasmid DNA was subjected to heat inactivation at 80 °C for 20 minutes. Plasmid DNA digests were run on 1% agarose gel at 105V for 50 minutes to determine successful digestion (**Fig. 2E**) before continuing to natural competence experiments with *F. nucleatum* 23726.

### Natural Competence Assays

A single colony was used to start overnight cultures of *F. nucleatum*. Stationary phase cultures were back diluted to OD_600_= 0.1 in 3 mL of Columbia Broth supplemented with hemin and vitamin K (CBHK) and grown to exponential phase (OD_600_= 0.5). 200 μL of cells were then transferred to 1.7mL microcentrifuge tubes. Unless indicated otherwise, 5 μg of DNA (genomic or plasmid) was added to the tubes, and the cultures were incubated for 4 hours at 37 °C under anaerobic conditions (90% N_2_, 5% CO_2_, 5% H_2_). The number of transformants was determined through plating on CBHK plates supplemented with 5 μg/mL thiamphenicol. Colony counts were analyzed after 48 hours of growth on selection media. As labeled on the Y-axis of **Figure 2**, we report the number of transformants per microgram of DNA from each 200 μL reaction.

### Statistics

All statistical analyses were performed in GraphPad Prism version 9. For single analysis, one-way analysis of variance (ANOVA) was used. For grouped analyses, two-way analysis of variance (ANOVA) was used. In each case, the following P values correspond to symbols in figures: ns (not significant), P > 0.05, *, P < 0.05; **, P < 0.01; ***, P < 0.001; ****, P < 0.0001. To obtain statistics, all studies were performed as three independent biological experiments. For all experiments in which statistical analysis was applied, an n= 3 of independent experiments was used.

## Data Availability

Data is available upon reasonable request.

## Acknowledgements

We would like to thank the Fralin Life Science Institute at Virginia Tech (Slade), and the USDA National Institute of Food and Agriculture (Slade) for funding. We thank the following individuals for help and guidance with these studies: Dr. Vic DiRita for critical insights into natural competency.

## Author Contributions

Blake E. Sanders; data curation, methodology, formal analysis, writing-original draft, review and editing: Ariana Umana; data curation, methodology, formal analysis, writing-original draft, review and editing: Tam T.D. Nguyen; data curation, methodology, formal analysis, review and editing: Kevin J. Williams; data curation, writing-review and editing: Christopher C. Yoo: data curation, writing-review and editing: Michael A. Casasanta; data curation, writing-review and editing: Bryce Wozniak; data curation, writing-review and editing: Daniel J. Slade: conceptualization, data curation, formal analysis, supervision, funding acquisition, validation, methodology, project administration, writing-original draft, review and editing.

## Supplemental Material

Available online as a dedicated file.

