## Supplemental Material for "Type 4 pili mediated natural competence in *Fusobacterium nucleatum*"

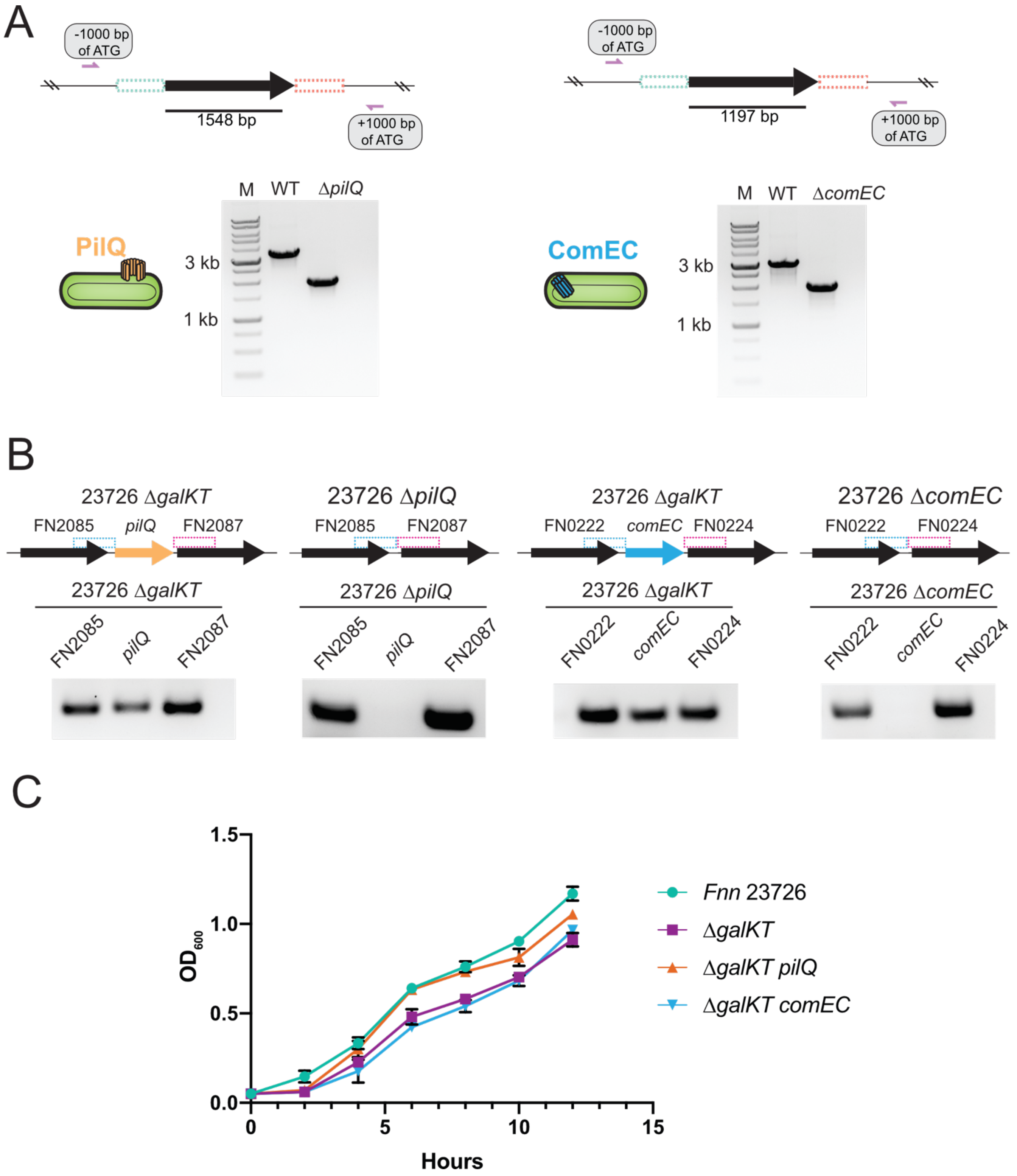


**Figure S1. Creation and validation of *F. nucleatum* 23726 Δ*galKT pilQ* and Δ *galKT comEC* (A)** Overview of primers constructed for confirmation of scarless *gspD and comEC* knockout mutant and PCR showing the markerless gene excision **(B)** Reverse-transcriptase PCR results verifying deletion of genes and absence of upstream and downstream polar effects **(C)** *F. nucleatum* 23726, Δ*galKT*, Δ*galKT pilQ, and* Δ*galKT comEC* strains were grown in

CBHK media and optical cell density was measured every two hours for a total of 12 hours to show that mutations do not affect growth.

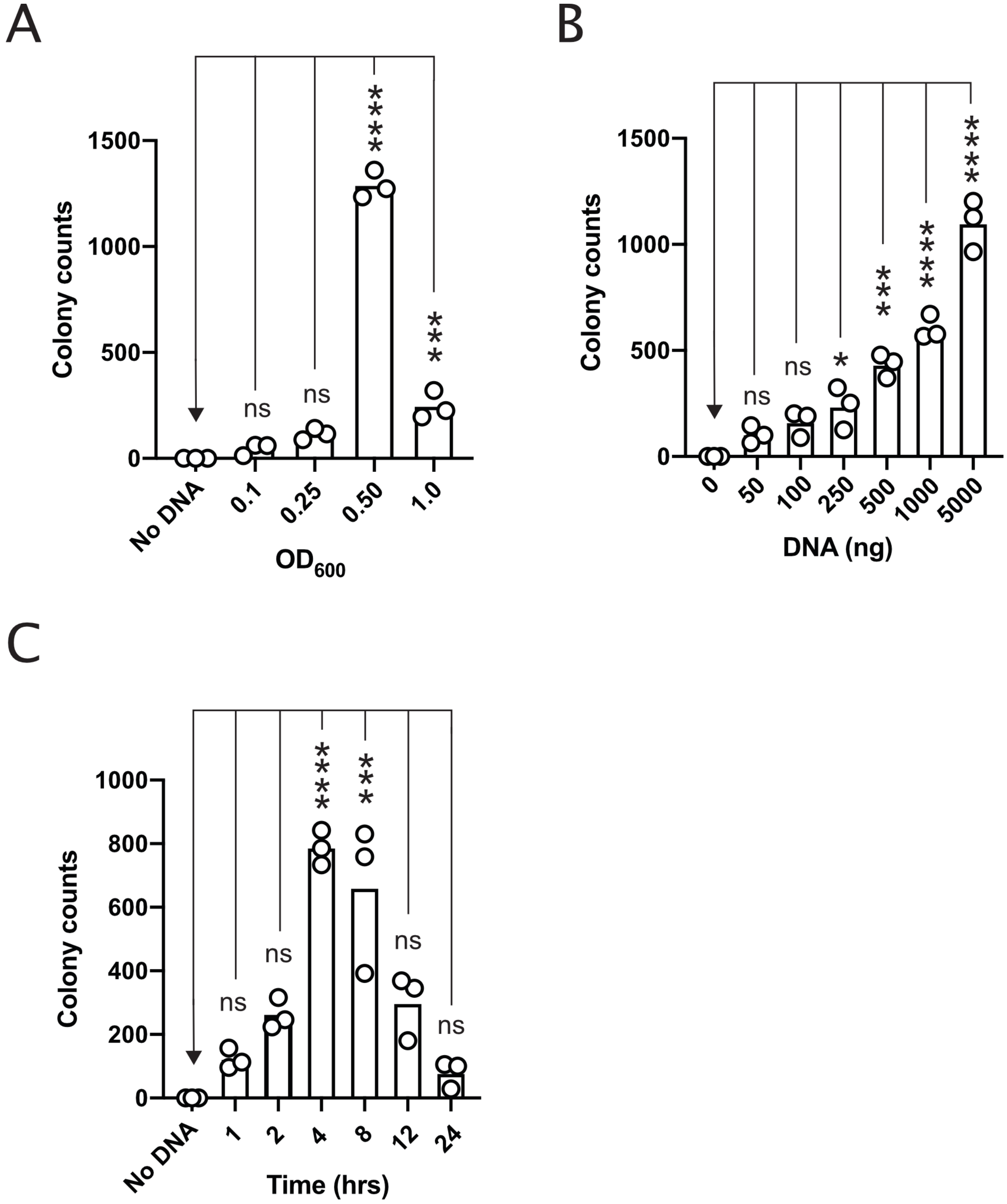


**Figure S2. Optimization of natural competence parameters in *F. nucleatum* 23726 for plasmid pDJSVT13. (A)** Incubation of 5 ug of methylated (M.Fnn23I, M.Fnn23II) pDJSVT13 for four hours at the indicated $OD_{600}$ readings. **(B)** Incubation of varying amounts of plasmid DNA at $OD_{600}$=0.5 for four hours. **(C)** Incubation of 5 ug of plasmid DNA at $OD_{600}$=0.5 for the noted times. Statistical values are as follows: ns (not significant), P.0.05;*, P,0.05;**, P,0.01;***, P,0.001;****, P,0.0001. Two-way ANOVA was used for panels A-C.

**Table S1. Competence genes present in *F. nucleatum* subsp. *nucleatum* ATCC 23726.**

| Gene Name | GenBank Protein ID^ | FusoPortal ID# | Amino Acids | Function |
|---|---|---|---|---|
| *comEA* | AVQ23988.1 : 159$ | 23726_Gene_112 | 176 | DNA pulling and import |
| *comEC* | AVQ24086.1 : 378$ | 23726_Gene_843 | 392 | DNA translocation through IM |
| *comF* | AVQ24217.1 : 204$ | 23726_Gene_1993 | 215 | DNA translocation through IM |
| *dprA* | AVQ23299.1 | 23726_Gene_1307 | 284 | ssDNA recombination |
| *pilA* | AVQ22554.1 | 23726_Gene_397 | 158 | Competence pilus (Major pilin) |
| *pilB* | AVQ22556.1 | 23726_Gene_399 | 414 | ATPase |
| *pilC* | AVQ22555.1 | 23726_Gene_398 | 346 | Competence pilus |
| *pilD* | AVQ22553.1 | 23726_Gene_396 | 165 | Competence pilus |
| *pilQ* | AVQ22550.1 | 23726_Gene_390 | 515 | OM Secretin |
| *pilT* | AVQ22319.1 | 23726_Gene_114 | 316 | ATPase |
| *pilV* | AVQ24018.1 : 128$ | 23726_Gene_395 | 137 | Competence pilus (Minor pilin) |
| *priA* | AVQ23220.1 | 23726_Gene_1214 | 766 | ssDNA recombination |
| *recA* | AVQ23748.1 | 23726_Gene_ 1846 | 378 | ssDNA recombination |
| *ssB* | AVQ23095.1 | 23726_Gene_1056 | 154 | ssDNA recombination |

^ GenBank Accession CP028109.1 and BioProject PRJNA433545

$ Incorrectly annotated by the NCBI in Genbank. Number of amino acids listed is less than that reported in Fusoportal, which in each case shows a correction in the N-terminus of the protein which contains the signal sequence critical to deliver the protein through the inner membrane via the Sec apparatus.

### Fusoportal.org[1]

**Table S2. Bacterial Strains used in this study.**

| Strain | Bacterial Species | Genotype and Characteristics | Reference |
|---|---|---|---|
| Top10 | *E. coli* | *mcrA* , Δ(*mrr-hsdRMS-mcrBC*), Phi80(del)M15, Δ*lacX74*, *deoR*, *recA1*, *araD139*, Δ(ara-leu)769, *galU*, *galK*, *rpsL*(SmR), *endA1*, *nupG* | Invitrogen |
| ArcticExpress (DE3) RIL | *E. coli* | B $F^-$ *ompT hsdS*($r_8^-$ $m_8^-$) $dcm^+$ $Tet^r$ *gal* λ(DE3) *endA* Hte [*cpn10 cpn60* $Gent^r$] [*argU ileY leuW* $Str^r$] | Agilent |
| LOBSTR-BL21(DE3)-RIL | *E. coli* | F–ompT hsdSB (rB- mB-) dcm gal (DE3) | [2] |
| *F. nucleatum* subsp. *nucleatum* ATCC 23726 | *F. nucleatum* | Wild Type | ATCC |
| *F. nucleatum* subsp. *nucleatum* ATCC 25586 | *F. nucleatum* | Wild Type | ATCC |
| DJSVT02 | *F. nucleatum* | *Fnn* 23726 ΔgalKT. In-frame deletion of *galK* and *galT* (Base strain for all target in-frame gene deletions) | [3] |
| BESVT1 | *F. nucleatum* | *Fnn* 23726 Δ*galKT* Δ*pilQ.* In-frame deletion of *pilQ* in the DJSVT02 background | This study |
| BESVT2 | *F. nucleatum* | *Fnn* 23726 Δ*galKT* Δ*comEC.* In-frame deletion of *comEC* in the DJSVT02 background | This study |

**Table S3. Primers used in this study.**

| Primer Name | Sequence (5' to 3') | Description |
|---|---|---|
| prBES01 | gcactaGGTACCCTTGATATAAAAAGAAAGGATTTTGAAAAGGATAGAGAAAAAATTG | Forward primer -750 bp upstream of *pilQ* in *F. nuc* 23726. Has a KpnI site. Makes construct pBES1. |
| prBES02 | TGAATTTACCATATTTTTTAAAAAATTTTTCTGTTCTCACCTTTTCTTAATTATATATAAGTCTACTTGACAG | Reverse primer -1 bp upstream of *pilQ* in *F. nuc* 23726. Overlaps with prBES03 for OLE-PCR. Makes construct pBES1. |
| prBES03 | AAAAATTTTTTAAAAAATATGGTAAATTCATTTTACATTCATTTTTCTACTATATAC | Forward primer +1 bp downstream of *pilQ* in *F. nuc* 23726. Overlaps with prBES02 for OLE-PCR. Makes construct pBES1. |
| prBES04 | gcactaACGCGTGATTATTCACTGAATTAGCATTTTTATTCCCTTTTTGTTC | Reverse primer +750 bp downstream of *pilQ* in *F. nuc* 23726. Has a MluI site. Makes construct pBES1. |
| prBES05 | CTATATTTTGAGAAAAGAAAATGAAATTGAATTTAAACTTGAAATTAAATACTTTTAG | Forward confirmation primer -1000 bp upstream of *pilQ* in *F. nuc* 23726. |
| prBES06 | GAAACTTGTGGCTTTTAATCTATATTAACTTTATTAGGGTG | Reverse confirmation primer +1000 bp downstream of *pilQ* in *F. nuc* 23726. |
| prBES07 | GATTGAAAGTAAGACTACAACTGAAAATAAAGAAGACAAG | Forward primer -250 bp *pilQ* *F. nuc* 23726 start for sequencing. |
| prBES08 | CTTTCTTAATCTCTCGTACATTTAGTATATATAATTAAAATGAATACCTAATG | Reverse primer +250 bp *pilQ* *F. nuc* 23726 start for sequencing. |
| prDJSVT1244 | gagctagaggtaccGGAAAACGAGAAATCACATTGGTATCAG | Forward primer -750 bp upstream of *comEC* in *F. nuc* 23726. Has a KpnI site. Makes construct pBES2. |
| prDJSVT1245 | CAATATAAATTAAATATTTTCAAAATAAAGATAGTAGTATTATAATCCTAAGAACTCACCTATATATTATCTTTTTTATATAG | Reverse primer -1 bp upstream of *comEC* in *F. nuc* 23726. Overlaps with prDJSVT1246 for OLE-PCR. Makes construct pBES2. |
| prDJSVT1246 | CTATATAAAAAAGATAATATATAGGTGAGTTCTTAGGATTATAATACTACTATCTTTATTTTGAAAATATTTAATTTATATTG | Forward primer +1 bp downstream of *comEC* in *F. nuc* 23726. Overlaps with prDJSVT1245 for OLE-PCR. Makes construct pBES2. |

| | | |
|---|---|---|
| prDJSVT1247 | cgtgatcgtacgcgtGCTGTTGCTGCATTACGAAGTTTATCAATTTC | Reverse primer +750 bp downstream of *comEC* in *F. nuc* 23726. Has a MluI site. Makes construct pBES2. |
| prDJSVT1248 | GTACTTTTTGGAATTAGAGGAATGATAATATTTTTATG | Forward confirmation primer -1000 bp upstream of *comEC* in *F. nuc* 23726. |
| prDJSVT1249 | CTAGTCTATATCCATTACTCATATATGATG | Reverse confirmation primer +1000 bp downstream of *comEC* in *F. nuc* 23726. |
| prDJSVT1250 | CAGATAGCAAGAAAGCTATTATAAAGAAATAAC | Forward primer -250 bp *comEC F. nuc* 23726 start for sequencing. |
| prDJSVT1251 | GCTCCTCCATTAAAATTGATTGTCTC | Reverse primer +250 bp *comEC F. nuc* 23726 start for sequencing. |
| **RT-PCR primers for gene KO confirmation** | | |
| prBES09 | GAATTTGATAATATTCCTAACTTATTTAACTATATAGAAGAAAAGATTTC | Forward primer for RT-PCR of FN2085, gene upstream of *pilQ* |
| prBES10 | GTTGTTTTTACTTCTAGTAAATTATTATTTACACTAATTTTTAGATATGAG | Reverse primer for RT-PCR of FN2085, gene upstream of *pilQ* |
| prBES11 | GAAACTTTTGGGGAGAATATAAAGGTAAGTAC | Forward primer for RT-PCR of FN2086, *pilQ* |
| prBES12 | CTAAAATCATTGCTGTCAGACTATTTCTTTC | Reverse primer for RT-PCR of FN2086, *pilQ* |
| prBES13 | GCTGAATTAAATGAACAACAAGCAAAAG | Forward primer for RT-PCR of FN2087, gene downstream of *pilQ* |
| prBES14 | GGCTTTATCAGAAGAAATTTTTGGATTATTCAC | Reverse primer for RT-PCR of FN2087, gene downstream of *pilQ* |
| prBES55 | GTGAGTTTTAAAGATACAATAGGAATTATAATAAA | Forward primer for RT-PCR of FN0222, gene upstream of *comEC* |
| prBES56 | CATAACATAATCTTTCTATAACCATATTTTTTAGA | Reverse primer for RT-PCR of FN0222, gene upstream of *comEC* |
| prBES57 | GGTCATTGCTATTTTTTATTTTATCTTTAAGA | Forward primer for RT-PCR of FN0223, *comEC* |

| | | |
|---|---|---|
| prBES58 | CGATATAGAAAAAATAACTGTTGGATTAATAAA | Reverse primer for RT-PCR of FN0223, *comEC* |
| prBES59 | GATAGCATAGTAAAAAATATAGAAAATGGAGT | Forward primer for RT-PCR of FN0224, gene downstream of *comEC* |
| prBES60 | GTTGTCTTTATATAAGCTTCTGGTTG | Reverse primer for RT-PCR of FN0224, gene downstream of *comEC* |

**Table S4. Plasmids utilized in this study**

| Plasmid name | Description | Reference |
|---|---|---|
| pDJSVT1 | Vector containing homologous regions +/- 1000 bp upstream and downstream of *galKT* for single crossover Integration in *F. nucleatum* 23726 and 25586 ($Cm^r$, $Tm^r$) | [3] |
| pDJSVT14 | pDJSVT13 with all CATG sites removed by OLE-PCR building of the vector ($Cm^r$, $Tm^r$) | [4] |
| pDJSVT24 | pET16b vector containing M.FnuI with constitutively active Anderson medium promoter ($Amp^r$) | [4] |
| pDJSVT25 | pET16b vector containing M.FnuII with constitutively active Anderson medium promoter ($Amp^r$) | [4] |
| pDJSVT26 | pET16b vector containing M.FnuI and M.FnuII with constitutively active Anderson medium promoter ($Amp^r$) | [4] |
| pBES1 | *pilQ* gene deletion vector for *F. nuc* 23726 ($Cm^r$ $Tm^r$) | This study |
| pBES2 | *comEC* gene deletion vector for *F. nuc* 23726 ($Cm^r$ $Tm^r$) | This study |

$Cm^r$, Chloramphenicol resistance
$Tm^r$, Thiamphenicol resistance
$Amp^r$, Ampicillin resistance
